# Viscoelasticity of basal plasma membranes and cortices derived from MDCK II cells

**DOI:** 10.1101/2021.05.20.444971

**Authors:** Andreas Janshoff

**Affiliations:** Department of Chemistry, Institute of Physical Chemistry, Göttingen

**Keywords:** Basolateral membrane, Actin cortex, Power law rheology, Pore-spanning membrane

## Abstract

In mature epithelial cells, however, cells adhere to one another through tight junctions, adherens junctions and desmosomes thereby displaying a pronounced apical-basal polarity. In vivo, the apical membrane has a larger surface area and faces the outer surface of the body or the lumen of internal cavities, whereas the basolateral membrane is oriented on the side away from the lumen and forms focal adhesions with the extracellular matrix. The mechanical properties of cells are largely determined by the architecture and dynamics of their viscoelastic cortex, which consists of a contractile, cross-linked actin mesh attached to the plasma membrane via linker proteins. Measuring the mechanical properties of adherent, polarized epithelial cells is usually limited to the upper, i.e., apical side of the cells due to their accessibility on culture dishes. Moreover, contributions from the cell interior comprising various filament types, organelles, and the crowded cytoplasm usually impede examination of the cortex alone. Here, we investigate the viscoelastic properties of basolateral membranes derived from polarized MDCK II epithelia in response to external deformation and compare them to living cells probed at the apical side. Therefore, we grew MDCK II cells on porous surfaces to confluency and removed the upper cell body by sandwich cleavage. The free-standing, defoliated cortices were subject to force indentation and relaxation experiments permitting a precise assessment of cortical viscoelasticity. A new theoretical framework to describe the force cycles is developed and applied to obtain the time-dependent area compressibility modulus of cell cortices from adherent cells. Compared to the viscoelastic response of living cells the basolateral membranes are substantially less fluid and stiffer but obey to the same universal scaling law if excess area is taken into account.

## Introduction

Cellular polarity is manifested at various levels such as the distribution of organelles, the composition of the plasma membrane and the architecture of the cytoskeleton.^1^ Particularly, epithelial cells exhibit polarized formation of cell-cell junctions comprising adherens junctions and tight junction separating the apical domain from the basolateral side.^2,3^ Many epithelial cells form microvilli at the apical domain filled with bundled actin filaments that increase the surface area substantially. On the basal side, stress fibers and focal adhesions emerge, responsible for attaching cells to the extracellular matrix. As for confluent polar epithelial cells, our understanding of cell mechanics comes mainly from indentation experiments probing the apical side facing the culture medium,^4–12^ whereas very few studies have addressed the elastic properties of the basal side let alone dissipative properties.^13–15^ It is generally believed that the response of cells to deformation originates predominately from the cellular cortex consisting of a thin, contractile and transiently cross-linked actin mesh connected to the plasma membrane.^16–19^ Pre-stress of the cortex is provided by motor proteins that in conjunction with cross-linked actin ensure resilience of the cell body on the one hand, and fluidity of the cell to perform dynamic shape changes, on the other hand.^17–21^ It was found that cells generally behave as a soft glassy material with power law exponents between 0.2 and 0.4 due to the broad distribution of relaxation times.^8,12,22–27^ Many weak interactions are involved in structure formation independent of molecular details, rendering the viscoelastic behavior appear unbound to a specific time scale.

Recently, we found that a linear viscoelastic continuum model of the cortex based on power law rheology explains consistently the viscoelastic response of nonpolarized MDCK II cells, confluent cells as well as apical cell membrane fragments over a wide time range.^19,28^ In particular, the defoliation of apical membranes by sandwich cleavage from living cells allowed to examine the impact of motor activity on fluidity.^28^ Based on these experiments and data from of a previous publications, I now also revisited indentation-relaxation experiments of basolateral membranes from MDCK II cells grown on porous supports.^5,13^ I first present a comprehensive theoretical model based on free-energy minimization that describes the viscoelastic response of thin membranes to indentation with a conical indenter and in a second step compared the stiffness-fluidity relationship to that of living cells probed from the apical side.

Compared to the viscoelastic properties of confluent cells, isolated basal cortices are much stiffer and less fluid, but follow the same universal scaling law when the excess area is properly accounted for.

## Theory

Here, I describe how force cycle experiments carried out with a conical indenter can be modeled with a minimum of essential assumptions to correctly access the viscoelastic properties of thin pore-spanning cortices. This comprises a scaling factor, the area compressibility modulus 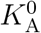, the pre-stress *σ*_0_, and the fluidity *β* (power law exponent) of the membrane-cortex composite. The general geometry is schematically shown in Fig. 1A. *R* denotes the radius of the pore and *a* is the contact radius of the membrane with the conical indenter.

**Figure 1:**
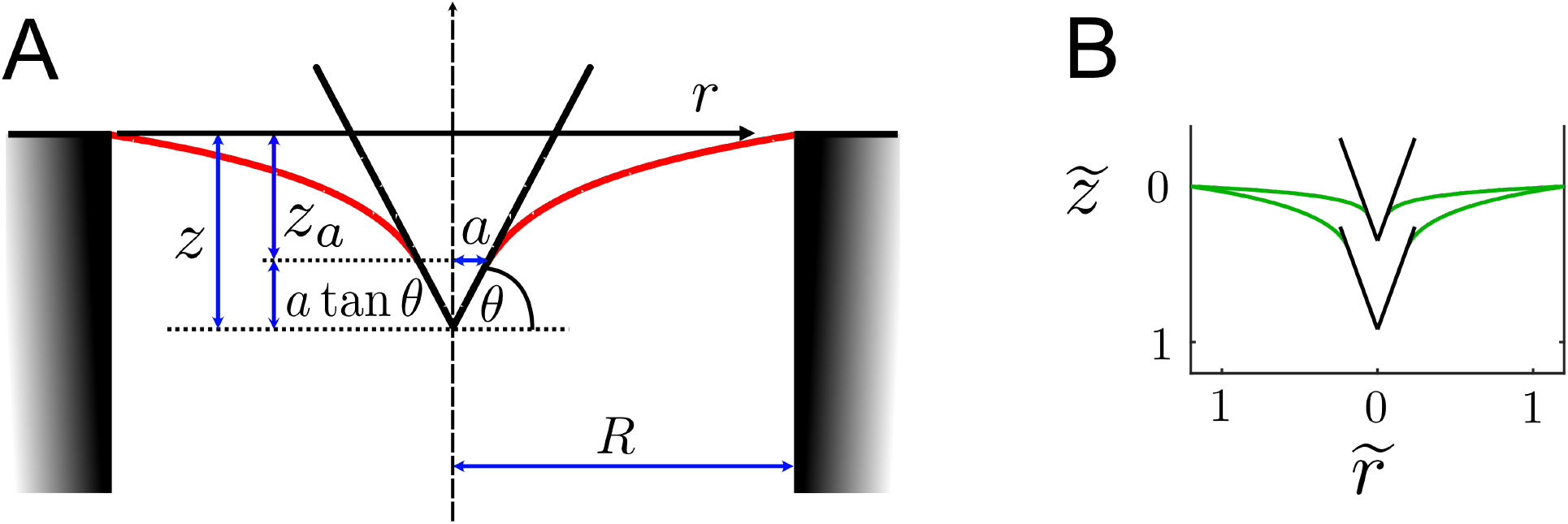
**A** Parametrization of the indentation experiment. The free membrane is in red color and the indenter in black. **B** General shape of the membrane (green) at two different indentation depths after minimizing the area.

### Force response of viscoelastic cortices spanned over a pore

In contrast to our previous publication that simplified the theoretical treatment by assuming that the AFM-indenter can be modelled by a cylindrical flat punch with a small radius of a few nanometers,^28^ I now consider a conical indenter and also abandoned the small gradient approximation, which is typically used to simplify the description of the force response.^9,29–32^ The following treatment is partly inspired by the work of Powers et al.^33^ who dealt with the formation of tethers pulled from a planar membrane. The free energy of the membrane is given by^34,35^

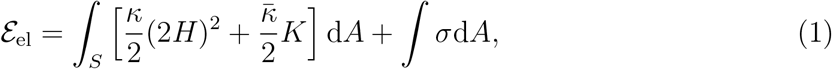

where *S* is the surface of the membrane, *H* its mean curvature (2*H* = 1/*R*_1_ + 1/*R*_2_) and K the Gaussian curvature (*K* = (*R*_1_*R*_2_^-1^). *κ* and 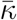 are the splay and saddle splay moduli, respectively, i.e., essentially elastic constants. While κ can easily be obtained experimentally and is always positive, 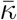 is difficult to obtain. The Gaussian curvature is independent of how the surface is embedded in ℝ^3^ and is an intrinsic property of the surface. According to the Gauss-Bonnet theorem the integral of the Gaussian curvature over a surface depends only on its topology and boundary. This implies that for a closed surface the energy contribution of the Gaussian curvature during any deformation is constant unless the topology of the surface changes and can be ignored when determining the shape of the membrane. Since the membrane has edges, the Gaussian modulus affects the shape through the boundary conditions, which we will neglect for the sake of simplicity, i.e., we set 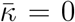.^33^ *σ* denotes the surface tension, i.e., comprising mainly the free energy contribution (per unit area) arising from adhesion of the cortex fragment to the pore rim, and in more general terms it represents the chemical potential of the membrane reservoir. I can also be considered a the Lagrangian multiplier to keep the area constant. The shape equation is obtained from standard variational calculus representing the balance of normal forces per unit area:^36^

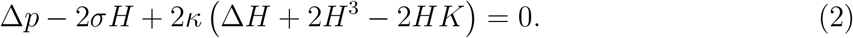

The pressure difference Δ*p* enters as the Lagrange multiplier ensuring constant volume. For free spanning membranes on pores open to both sides we can discard this contribution for the free membrane. We assume that the energy contribution due to recruiting new surface area against the surface tension *σ* ≈ 10^-3^ N is substantially larger then the bending energy. Since *κ* is expected to be rather small, on the order of 10^-19^ J depending on the thickness *d* of the cortex *κ* ∝ *d*^3^ and the lipid composition, we can assume that the dimensionless perturbation parameter *ε* = *κ*/*σ* is indeed very small (≈ 10^-16^m^2^). Thermal fluctuations are therefore negligible and we can rewrite equation (2):

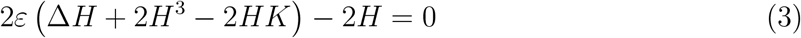

Assuming that *ε* = 0 we obtain *H* = 0, the minimal surface equation, which is essentially a catenoid since it is the only non-axisymmetric and non-planar minimal surface with zero mean curvature. It is, however, immediately clear that this simplified differential equation is not entirely compatible with the boundary condition since *ε* is multiplied with the highest derivative of *H*. Precisely, an external force is balanced by curvature where the indenter meets the membrane *r* = *a* implying that *H* ≠ 0. In contrast, the boundary condition σ*H* = 0 at *r* = *R*, the pore rim, is compatible with the differential equation as the pore rim acts as a hinge. Employing the concept of perturbation theory, we therefore need to consider a boundary layer at the contact line with the indenter rendering outer *r* > *δ* and inner (*r* < *δ*) solutions incompatible. The outer solution of the free membrane is a catenoid held between two circular boundaries, one being the pore with a large radius *R* and the smaller one defined by the contact with the conical indenter at *r* = *a*. The thickness of the boundary layer 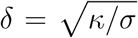 can be inferred from equating *ε*Δ*H* with *H* and represents a characteristic length scale. On scales larger than *δ* tension dominates, while on smaller length scales bending is the most important energy contribution. In our case, *δ* is on the order of 10 nm. In the following we will only consider the dominant outer solution since the characteristic length scale is governed by the surface tension due to the thin membrane patches and the large adhesion forces.

The problem of finding the shape *r*(*z*) of the membrane during indentation therefore reduces to the problem of finding its minimal free surface. We first consider the elementary case of two rings of equal size separating the membrane by 2*L* to form a shape with zero mean curvature. The area element d*A* = 2*πr*d*s* generates the surface through the integral

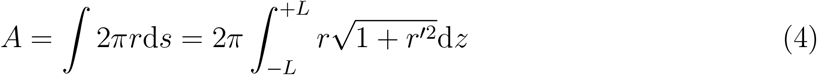

with *r*′ = *dr*/*dz*. Using standard techniques of variational calculus we arrive at

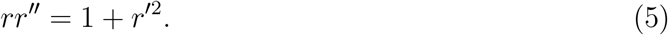

The differential equation can be integrated in two consecutive steps. Using (1 + *r*′^2^)′ = 2*r*′*r*′′ we obtain

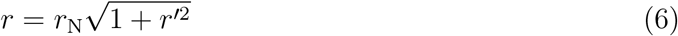

with *r*_N_ being a constant. Second, we employ the identity

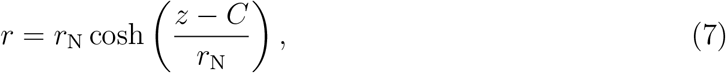

with the integration constant *C* being zero for two equally sized rings. Since we have one radius given by the pore rim *R* and the other one by the contact radius a we need to infer *C* from this boundary condition. *r*_N_ is identified as the minimal radius of the catenoid, its so-called neck radius. *C* can be obtained from *r* = *R* of the upper rim:

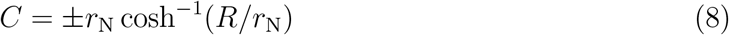

leading to

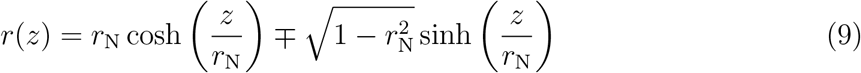

The upper sign corresponds to catenaries with a minimum neck radius at a positive value of *z*(*C* > 0), the indentation depth. Conversely, for *z*(*r*), the shape equation of the free membrane, we can write:

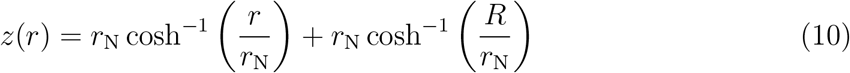

or in nondimensional form 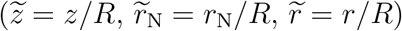:

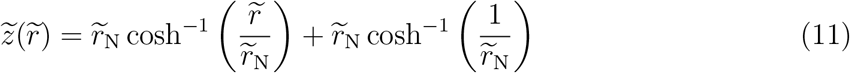

A simple relation holds between *r*_N_ the minimal radius and the force *f*:

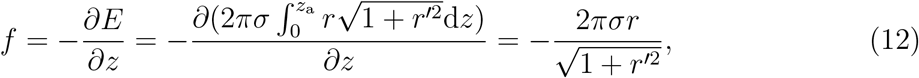

with *z_a_* the indentation depth at *r* = *a*. At the neck of the catenoid *r* = *K*, we have *r*′ = 0 and therefore *K* = *f*/(2*πσ*). Since at *r*(*z_a_*) = *a* with *a* > *K* we can write:

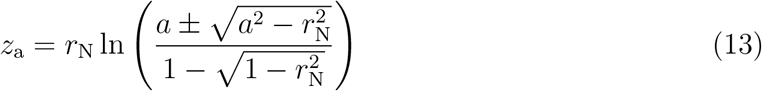

Equation 13 is responsible for two branches forming a closed curve in the *f* – *z_a_* plane for *a* < *R*. In principle, a critical (maximal) separation *z_a_* exists, where no solution is found, i.e. the catenoid becomes unstable and breaks. If the contact radius a is fixed and the indentation depth below the maximum (prior to instability) two catenoidal equilibrium solutions exist (see eq. 13). We only have to consider the branch (minus sign in eq. 13) with larger 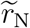 that has less area. The other branch is not found for real minimal surfaces. Therefore, we proceed with the minus sign in eq. 13. The existence of an elastic boundary layer allows the limit of a point force, i.e. *a* → 0, which is in contrast to pure soap films. In practice, however, point forces do not play a role since conventional AFM tips display curvature radii of approximately 20 nm. Now we only have to determine the contact radius *a* from the continuity condition, where the slope is identical for indenter and free-standing membrane. For a conical indenter we find:

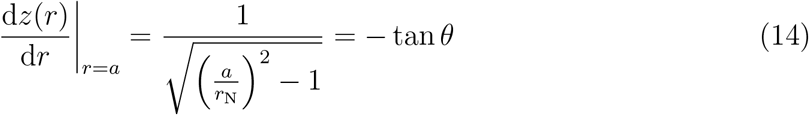

with *π*/2 – *θ* the half-opening angle of the cone giving

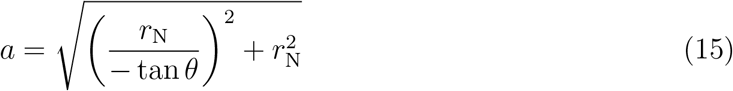

and the indentation depth at the tip of the indenter is *z*(*r* = 0) = *z_a_* + *a*tan(*θ*). Note that *a* > *r*_N_ as *r*_N_ is the smallest possible radius of a catenoid. The tension of the membrane is not necessarily a constant but depends on the area dilatation which inevitably occurs upon indentation.

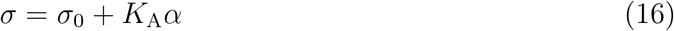

with *σ*_0_ the initial pre-stress and *K*_A_ the area compressibility modulus. 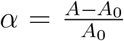 denotes the relative area dilatation with *A* the actual area and *A*_0_ the area prior to indentation, i.e., *A*_0_ = *πR*^2^. The actual area of the free membrane forming the catenoid is:

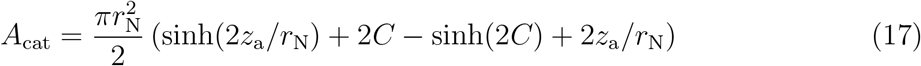

with *C* = – cosh^-1^(*R/r*_N_). Additionally, we need to consider coating of the cone up to *r* = *a* leading to *A*_cone_ = *πa*^2^/*cos*(*θ*) and therefore the overall area of the membrane is *A* = *A*_cone_ + *A*_cat_. If excess membrane area *A*_ex_ is recruited from the pore rim we refer to an apparent compressibility module 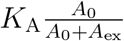.

As shown previously, viscoelasticity enters through the time dependency of the area compressibility modulus 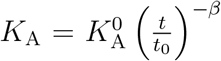 with 0 ≤ *β* ≤ 1 and *t*_0_ = 1s (set arbitrarily).^19^ The power law indicates that relaxation is not tied to an internal time scale.^22^ Consequently, the elastic-viscoelastic-correspondence principle leads to the following expression for the overall tension:

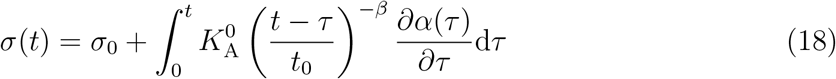

Since viscoelasticity of the membrane-cortex composite impacts only the in-plane area compressibility modulus, we can safely assume that the contour during indentation is identical to the contour for the elastic case. In nondimensional form 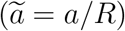 the indentation depth at *r* = 0 is^37^

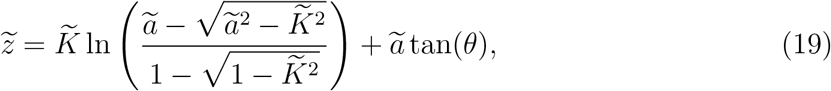

which tells us that for a given indenter geometry, i.e., *θ* value, the shape of the membrane and its scaled force response is uniquely defined by the distance between the two rings. The same is naturally true for the surface integral. This allows us to numerically compute 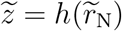 as well as 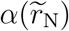 for each value of *θ* once and for all and fit the two curves with two polynomials, 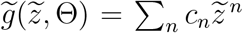 and 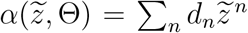, respectively. This permits us to obtain an analytical solution of the corresponding elastic-viscoelastic problem for indentation (approach)

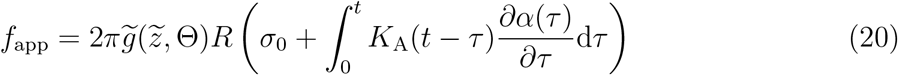

and retraction

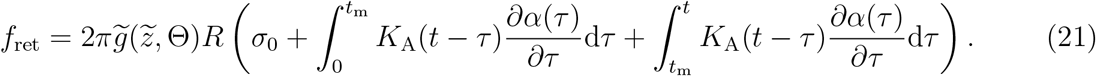

Here we assumed that in-plane stretching of the membrane/cortex is time dependent 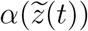 as we apply a linear ramp 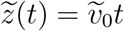 at the approach and 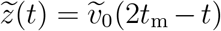 upon retraction, respectively 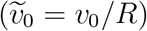. Hereditary integrals using a polynomial to the order *n* for *α*(*t*) are readily solved:

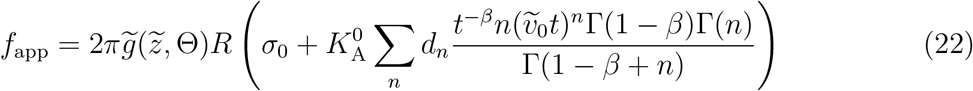

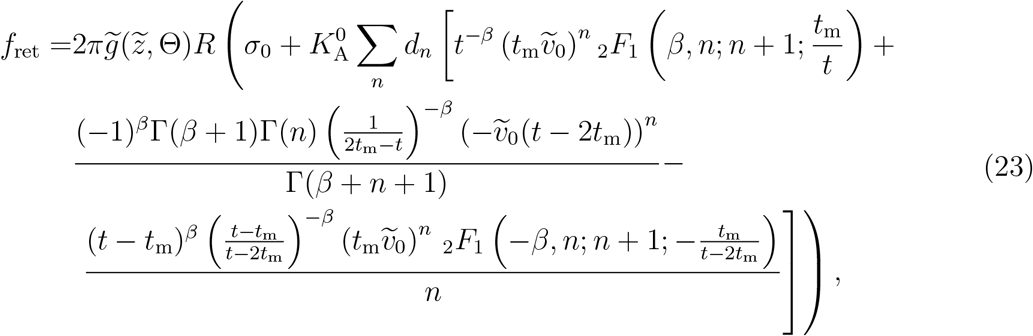

with the Gamma function 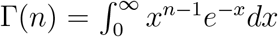 and the ordinary hypergeometric function 2*F*_1_(*a,b*; *c*; *z*). Usually, polynomials to the order of *n* = 4 are sufficient to describe the functions 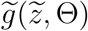 and 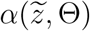 with sufficient accuracy. Experimental force - time curves were subject to fitting a piecewise function *f* (*t* ≤ *t*_m_) = *f*_app_(*t*) and *f*(*t* > *t*_m_) = *f*_ret_(*t*).

### Force response of living cells

We use the model von Hubrich et al. to fit the data^28^ In brief, confluent cells are as capped cylinders with contact angles around *ϕ*_0_; 35 ° prior to deformation.^38^ Generally, we consider the cell as a liquid-filled object surrounded by an isotropic viscoelastic shell deformed at constant volume. The force *f* acting on the apex of the cell:^28^

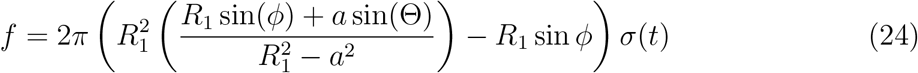

or in nondimensional form 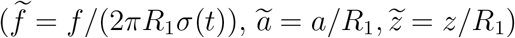

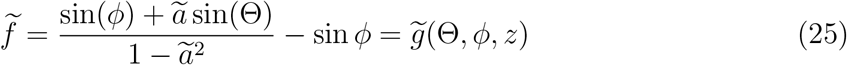

with *R*_1_, the radius at the base of the spherical cap and *ϕ* the contact angle in response to deformation.^5^ For a given set of angles *ϕ* and Θ the function 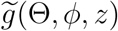 is computed numerically and the outcome fitted by by a polynomial 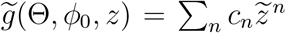 to obtain the coefficients *c_n_*. Computation of area change has been outlined before and the outcome approximated with a polynomial as described above 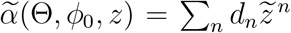 to determine *d_n_*, respectively.^5,28^

Viscoelasticity of the shell is included through equation (18) replacing *σ*(*t*). What follows is numerical solution of a set of non-linear equations for the shape of the deformed cell to fulfil force balances and the constant volume boundary condition.^28^ For a given indenter geometry and contact angle ϕ_0_ this has to be solved only once and scaling is accomplished by multiplication with *R*_1_.

## Results and Discussion

Previously, we investigated the topography and elastic properties of basolateral membranes derived from confluent MDCK II cells grown on porous substrates.^13^ Defoliation was accomplished according to the squirting - lysing protocol, in which the confluent MDCK II cells were first subject to osmotic swelling with addition of hypotonic buffer (Fig. 2A).^13,39^ The cells were ruptured by applying a gentle buffer stream from a syringe directed to the cell monolayer at an angle of 45 °. The indentation curves obtained from the pore’s center were previously described using an asymptotic linear relationship between force and indentation depth essentially capturing only the pre-stress of the cortex.^13^ I now reevaluated the data, including also the retraction curves that were not considered in the previous publication, by applying the viscoelastic model described above. Both indentation and relaxation were fitted with equations (22, 23) as a piecewise function providing access to three relevant mechanical parameters, the pre-stress *σ*_0_, the scaling factor (apparent area compressibility modulus) 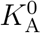 and the fluidity or power law exponent *β*, the latter two as we will see are not independent. It is important to notice that for pore-spanning membranes and cortices, the pre-stress *σ*_0_ corresponds mainly to the differential adhesion free energy between the pore rims *G*_pr_ and the free-standing part *G*_p_

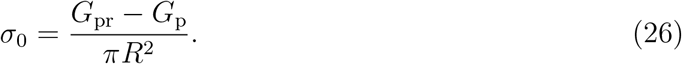

**Figure 2:**
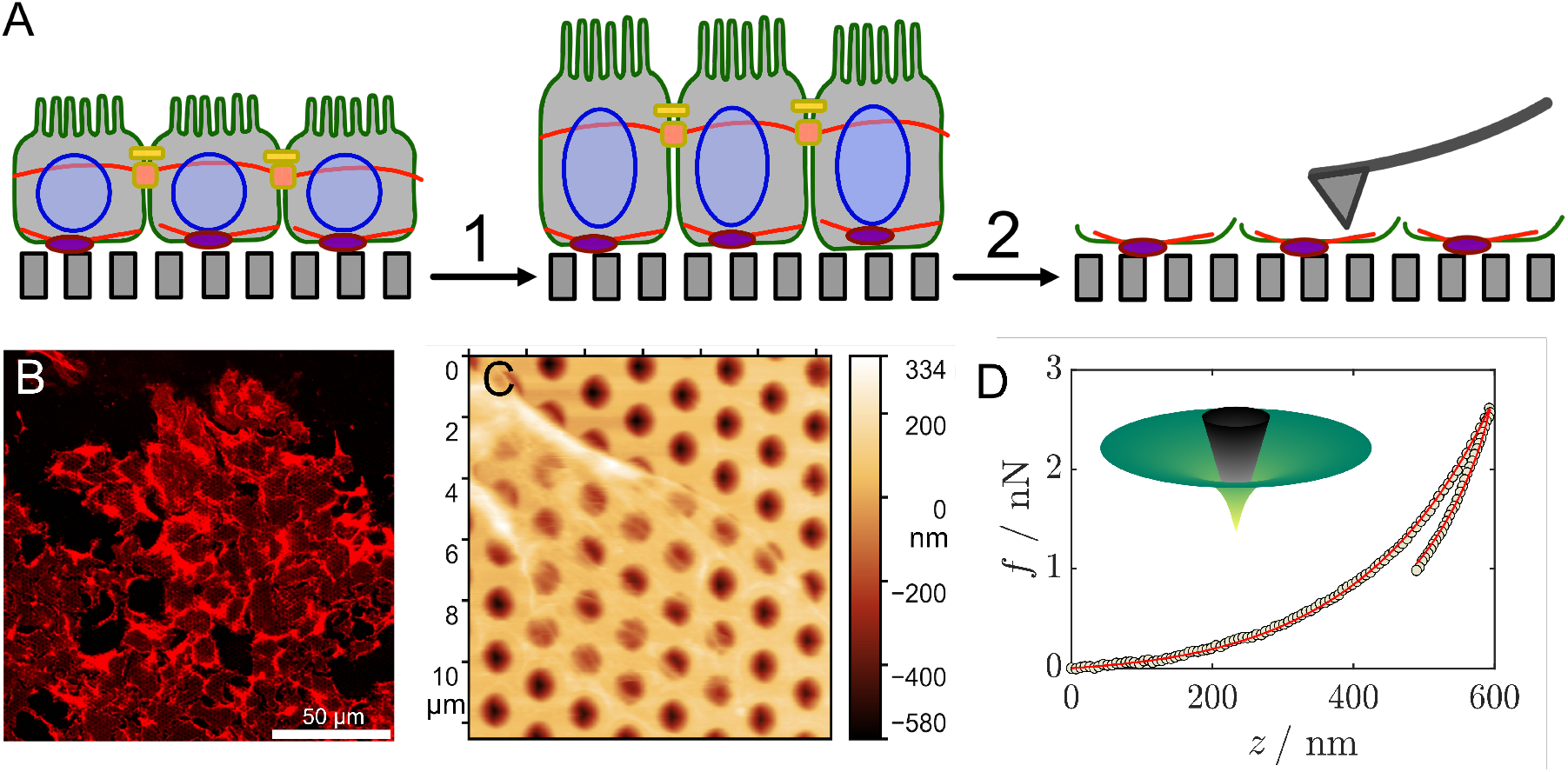
**A** Preparation of basolateral membrane sheets on porous supports.^13^ MDCK II cells are first grown to confluence on a porous support with pores of 1.2 *μ*m in diameter. After exposing the cells to hypotonic solution (1) the swelling makes them susceptible for shear stress (2) applied by a buffer stream. The isolated membranes were subject to indentation experiments using a conventional AFM instrument. **B** Fluorescence micrograph showing the results of the squirting-lysing protocol (TRITC - phalloidin stained actin filaments). **C** AFM image showing actin stress fibres. **D** Typical force indentation - relaxation curve (circles) subject to fitting of equations (22, 23) (red line). Fitting range was limited to avoid interference from adhesion events. The inset shows the shape of the membrane at largest indentation depth.

The area compressibility modulus is the response function of the linear - elastic resistance of the cortex/membrane assembly against in-plane area dilatation. Depending on the boundary conditions, area dilatation occurs inevitably during deformation as required for deviation from a minimal surface. While cells maintain a constant volume during deformation, the basal membrane sheets are physically and chemically attached to the pore rims - the strength of attachment given by equation (24).^19,40,41^ The measured area compressibility modulus of the basolateral membrane sheets contains contributions from the rather inextensible membrane and the actin mesh. Albeit the outstretched plasma membrane is almost inextensible exhibiting considerable large *K*_A_ values of 0.1 - 0.5 N/m depending on the lipid composition,^42^ excess membrane area *A*_ex_ can be recruited from wrinkles and folds during indentation. This diminishes the measured area compressibility by a factor of *A*_0_/(*A*_0_ + *A*_ex_). Experiments with neat lipid bilayers neither display a measurable area compressibility modulus nor a hysteresis during relaxation.^32^ The second contribution to *K*_A_ comes from the underlying actin cortex, which points towards the indenter in this case. Notably, recruitment of excess area from the adjacent surface is also possible in this case, giving rise to apparent values (*vide infra*), which are substantially smaller compared to the isolated pore. As pointed out previously, knowledge of cortex thickness and mesh size allows to roughly estimate the elastic modulus of a cross-linked actin network:^19,43^

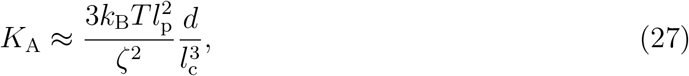

with the distance between cross-links 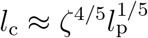 and the persistence length *l*_p_.^43^ Assuming reasonable values for the mesh size of *ζ* = 100 nm, a cortex thickness *d* of 150 nm and a persistence length of 17 *μ*m^44^ we obtain a *K*_A_ values of about 2.5 mN/m. In the previous publication we removed filamentous actin partly resulting in a substantial softening of the membrane patch.^13^

The power law exponent *β* represents the flow behaviour of the cortex. If *β* is close to zero, the cortex behaves as an elastic solid, whereas a *β* value of one corresponds to a Newtonian liquid. Generally, intermediate values are found for living cells. It could be shown that *β* is not independent of the corresponding elastic modulus or scaling factor in the case of power law rheology, which in our case is the apparent area compressibility modulus 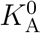, but decreases according to 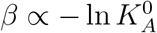 (*vide infra*).^19^ Fig. 2D shows a typical force cycle curve consisting of an approach curve generated by a linear ramp and a subsequent relaxation also following a linear ramp (identical approach and retraction velocity) obtained by probing a basolateral membrane patch. The patch covering the pore was indented precisely in the center fulfilling the axial symmetry conditions of the theory. Due to the potential for adhesion events to affect the retraction curve, only a portion of the retraction curve (approximately 1/3) was considered for fitting.

Fig. 3 shows the results of fitting equations (22, 23) to experimental force curves obtained from basolateral membrane patches of adherent MDCK II cells. The mean pre-stress 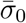 was 1.6 ± 1 mN/m, which we largely attribute to differential adhesion at the pore rim and stress exerted by actin bundles. The large variance is due to substantial pore to pore variation supporting the theory of differential adhesion that can locally be quite different, for instance in the vicinity of focal contacts. Most importantly, a linear decrease of *β* with ln 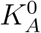 was found, which could be expected considering the reported scaling behavior of living cells and apical membrane fragments.^22,28,45^ Since *β* and 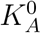 depend on each other in this way it implies automatically that stiffer cortices are also less fluid and vice versa. This universal law has first been discovered by Fabry et al. using ferrimagnetic beads adhering to the actin cytoskeleton and actuated by an external magnetic field.^22^ A weak power law was observed for *G*′ being larger than *G*” below 300 Hz. The spectra were described by a structural damping model that produced parameters falling onto a master curve. This essentially suggested that the constitutive elastic and frictional properties are controlled by a single parameter, for instance, *β* over a wide frequency range.^22^ Thus, in principle, the cells can respond to external cues by modulating solely *β* as the control parameter. However, if the data are extrapolated to a fully elastic material (*β* = 0, intersection with the *x*-axis) the basolateral membranes exhibit a very low stiffness compared to living cells (Fig. 3).^28^ This needs further exegesis. If one takes into account that the excess area might be as large as the patch itself including hidden reservoirs *A*_ex_ ≈ 329 ± 49 *μ*m^46^ since upon indentation the cortex can follow the indenter into the pore, the corrected (see arrow in Fg. 3) correlation (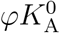 with 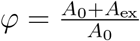) continues the scaling found for living cells probed at the apical side (Fig. 3). Therefore, considering only one parameter, the fluidity *β* is sufficient. The lack of functional motor proteins could be responsible for the flatter slope found for the basolateral membrane patches compared to that of living cells. This was shown recently for apical cell membrane fragment after addition of exogenous ATP to revive some of the remaining myosin motors significantly increasing fluidity of the cortex.^28^ A direct comparison between cortex fragments derived either from the basolateral or the apical side show that polarity has only a small impact on viscoelasticity. Hubrich et al.^28^ found that apical membrane fragments also exhibit low fluidity in the range of 0.2 similar to what was found here, while living cells display substantially higher fluidity presumably due to motor activity as mentioned above. Along the same lines, Kim et al. found only small differences in elasticity of PaTu8988S and PaTu8988T probed either from the basal or apical side, respectively.^15^

**Figure 3:**
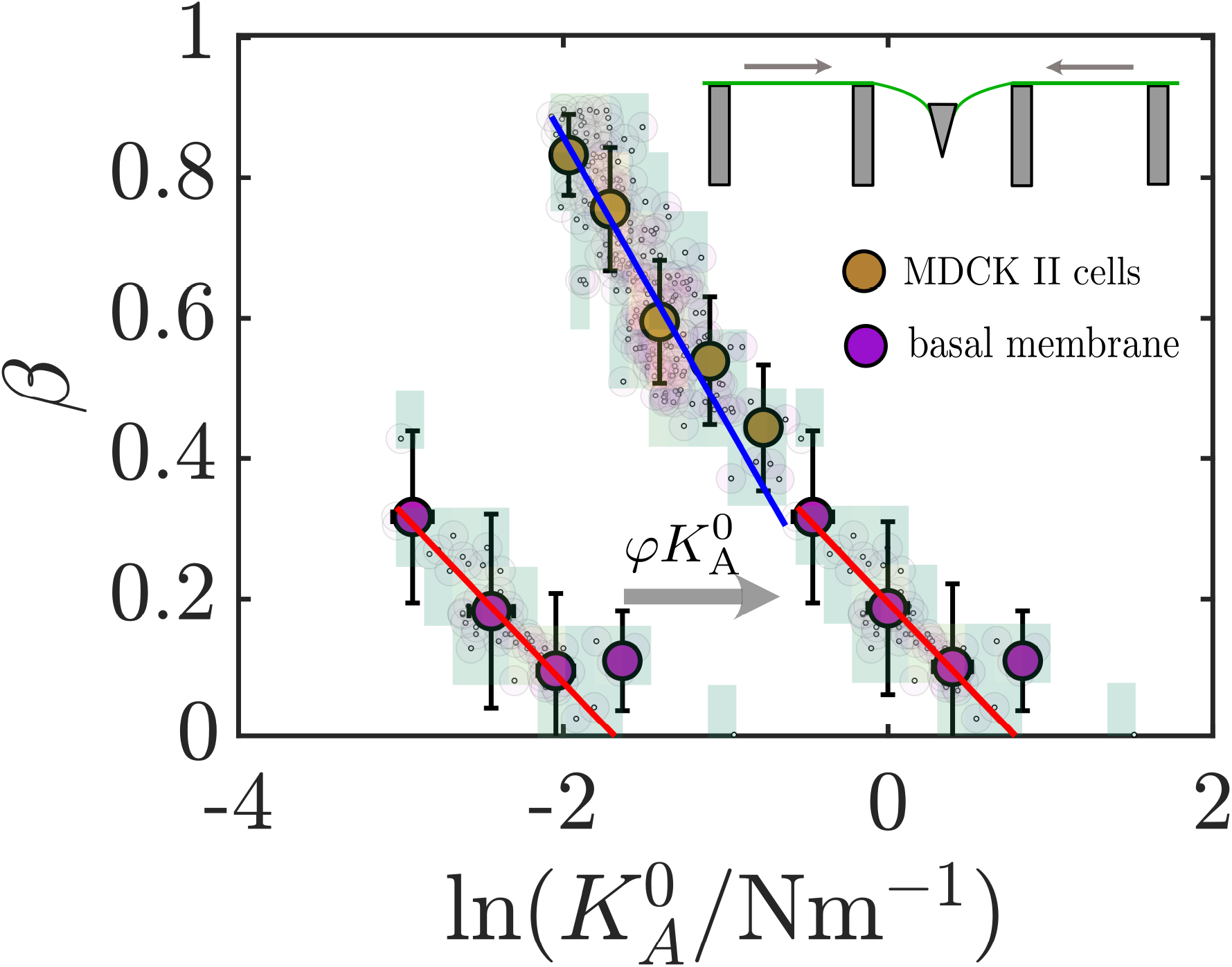
Fluidity *β* as a function of area compressibility 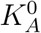 for basolateral membrane sheets (purple) and living cells probe from the apical side (dark yellow). The data for living cells were obtained from reevaluating experiments by Pietuch *et al.*^5^ using the model of Hubrich *et al.*^28^ The arrow illustrates what happens to the data if 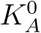 is rescaled by the average area of a membrane footprint, the flow illustrated by the inset in the top right corner.

Notably, the presence of pores during cell culture renders MDCK generally softer than cultured on continuous stiff surfaces such as culture dishes or silica. This was recently shown in a systematic fashion using different pore sizes.^47^ The area compressibility modulus might be reduced by a factor of 2-3 comparing cells cultured on flat substrates with those grown on porous surfaces with a pore diameter of 5 *μ*m.^47^

In conclusion, for the first time it was possible to obtain the viscoelastic properties of baso-lateral membranes in the absence of other cellular ingredients by site-specific indentation - relaxation experiments of planar membrane patches on porous substrates. The theoretical model to describe the force cycles correctly describes the shape of the free membrane/cortex in terms of a minimal surface and permits to easily modify the constitutive equations to capture dissipative processes in thin films employing the viscoelastic-elastic correspondence principle.^8^ It could be shown that the universal scaling law between stiffness of cells and their fluidity is largely preserved, implying that cell cortices cannot change their elastic and dissipative properties independently.^19,22,45^ A decrease of stiffness is always accompanied by an increase in fluidity. We found that regulation of mechanical properties can also be accomplished via storage of excess area to soften the apparent modules over orders of magnitude. The cortex fragments are stiffer and less fluid compared to living cells which can be partly attributed to arrested myosin motors but could also be a consequence of polarity and therefore larger pre-stress exerted by stress fibers at the basal side. While for unstressed actin networks a power law exponent of 0.5 is expected, we found substantially lower values of only 0.2.^27^ Missing motor activity decreases the noise level necessary to drive the cytoskeleton into a disordered state that eventually enables the cells to perform tasks like spreading, migration and division.

Polarity of epithelial cells is very pronounced on many levels and cortical viscoelasticity seems to be no exception. However, more experiments involving fully active cortices with myosin motors are needed to obtain a more comprehensive picture of viscoelasticity in the context of cell polarity.

## Material and Methods

All experimental data were obtained from previous publications. Experimental force data from confluent MDCK II were taken from Pietuch *et al.*,^5^ while all data from basolateral membrane patches were published by Lorenz *et al.*^13^ In these works only the approach curves were evaluated using exclusively elastic tension models. Here, I included the available retraction curves in the comprehensive viscoelastic analysis. The following paragraphs repeat the key steps in obtaining these data.

### Preparation of baslolateral membrane patches on porous substrates

Porous silicon substrates purchased from fluXXion B.V. were used as cell culture substrates. Pores possess a depth of 800 nm and display a diameter of 1.2 *μ*m. The substrates were first coated with a thin adhesive layer of chromium (3 nm) followed by a gold coating of 60 nm, which has been proven to be an excellent surface for culturing MDCK II cells.^48^ Basolateral membrane fragments of MDCK II cells were obtained using a squirting - lysing protocol described previously.^13,39^ Confluent MDCK II cells were grown in minimal essential medium (MEM) on porous substrates supplemented with 2 mM L-glutamine and 10% (*v/v*) fetal calf serum (FCS) at 37° in an 95% air/5% CO_2_ humidified incubator. Cell were subject to osmotic stress using hypotonic buffer. Shear stress by a buffer stream led to cleavage of the cells and also eventually to complete removal of the upper cell bodies.

### Atomic force microscopy and indentation experiments

AFM measurements were carried out with a MFP-3D microscope (Asylum Research, Santa Barbara, CA, USA) at 20° using MSCT cantilevers from Veeco with a nominal spring constant of 0.01 N/m. Ramp velocities were kept constant and set to 2 *μ*m/s.

## Acknowledgement

This work was supported by the VW foundation (Living Foams) and the DFG JA 963/18-1. I thank A. Pietuch, B. Lorenz and I. Mey for providing the experimental data and stimulating discussions.

## TOC Graphic

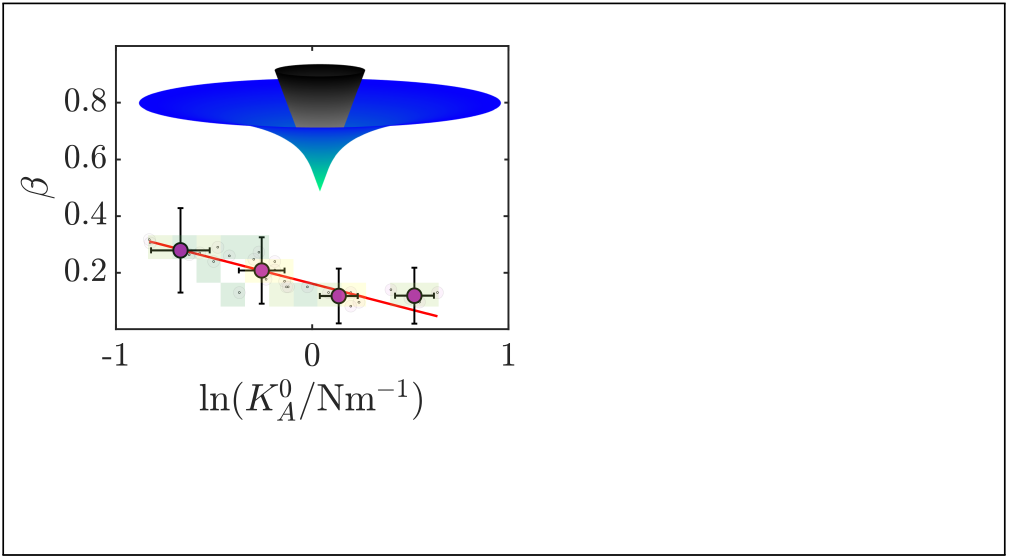

